# Memory encoding and retrieval by retrosplenial parvalbumin interneurons is impaired in Alzheimer’s disease model mice

**DOI:** 10.1101/2023.09.02.555835

**Authors:** Kyerl Park, Michael M. Kohl, Jeehyun Kwag

## Abstract

Memory deficits in Alzheimer’s disease (AD) show a strong link with GABAergic interneuron dysfunctions^1–7^. Ensemble dynamics of GABAergic interneurons are critical in memory encoding and retrieval^8–12^ but how GABAergic interneuron dysfunction affects inhibitory ensemble dynamics in AD is unknown. As retrosplenial cortex (RSC) is a brain area critical for episodic memory^13–16^ and affected by beta-amyloid accumulation in early AD^17–21^, we address this question by performing Ca2^+^ imaging in RSC parvalbumin-expressing (PV) interneurons during a contextual fear memory (CFM) task in healthy control mice and the 5XFAD mouse model of AD. We found that subpopulations of PV interneurons that were responsive to aversive electric foot shocks during contextual fear conditioning (CFC) in the control mice significantly decreased in the 5XFAD mice, indicating a dysfunction in the recruitment of CFM-encoding PV interneurons. In control mice, CFC-responsive PV interneuron ensemble activities were selectively upregulated during the freezing epoch of the CFM retrieval, manifested by CFC-induced synaptic potentiation of PV interneuron-mediated inhibition. However, CFC-induced changes in PV interneuron ensemble dynamics during CFM retrieval and synaptic plasticity were both absent in the 5XFAD mice. Optogenetic silencing of PV interneurons during CFC in control mice mimicked the CFM deficits in 5XFAD mice, while optogenetic activation of PV interneurons during CFC in the 5XFAD mice restored CFM retrieval. These results demonstrate the necessity and sufficiency of CFC-responsive PV interneurons for CFM retrieval and suggest that synaptic dysfunction in CFM-encoding PV interneurons disrupt the inhibitory ensemble dynamics underlying CFM retrieval, subsequently leading to memory deficits in AD.

## Results and Discussion

### Encoding of contextual fear memory by RSC PV interneurons is impaired in 5XFAD/PV-Cre mice

Neurons in the retrosplenial cortex (RSC) have been suggested to encode contextual fear memory (CFM)^14, 15^, but the contribution of GABAergic interneurons remains unclear. Among GABAergic interneurons, which have been reported to play a key role in memory functions^8–12^, parvalbumin-expressing (PV) interneurons are the most abundant GABAergic interneurons in the RSC^22^. Thus, we first set out to compare the ensemble dynamics of RSC PV interneurons during CFM encoding and retrieval between control mice and 5XFAD mouse model of AD. To do so, we injected AAV-DIO-GCaMP6s into the granular RSC of PV-Cre and in 5XFAD/PV-Cre mice (Figure 1A, left), from which longitudinal Ca^2+^ imaging of GCaMP6s-expressing PV interneurons was performed using a miniaturized microscope (Figure 1A, right) during CFM task (Figure 1B). In the CFM task, mice were first habituated to the test chamber (day (d) 1, habituation). A day later (d2), mice were placed in the same chamber for contextual fear conditioning (CFC), during which associative memory between the context of the chamber and the 1 second-long aversive electric foot shocks is formed^23^. On the third day (d3), mice were returned to the chamber for the testing of CFM retrieval measured as freezing behavior (Figure 1B, C). In 5XFAD/PV-Cre mice that showed beta-amyloid accumulations in the granular layers of RSC (Figure S1A), freezing behavior was significantly decreased during CFM retrieval session compared to PV-Cre mice, which was independent of locomotion of 5XFAD/PV-Cre mice (Figure S1B, C), confirming that CFM retrieval is impaired in 5XFAD/PV-Cre mice (Figure 1C). From the Ca^2+^ signals of neural ensembles acquired during the entire CFM task duration, we first analyzed how CFM encoding is represented in the Ca^2+^ dynamics of PV interneuron ensembles. Extraction of spikes by deconvolutions of Ca^2+^ signals (Figure 1D) during CFC session revealed that subpopulations of PV interneuron ensembles were responsive to aversive electric foot shocks during CFC, herein termed CFC-responsive (CFC-R) neurons, while the rest were not responsive to aversive electric foot shocks, herein termed CFC non-responsive (CFC-NR) neurons (Figure 1D-F). A PV interneuron was defined to be CFC-R if the spike firing rate in response to the electric foot shocks (CFC response) exceeded the 95th percentile of the probability distribution of randomly shuffled shock response while the remaining neurons were defined as CFC-NR neurons (Figure. 1E). Based on these analyses, in PV-Cre mice, 55% of PV interneurons (68 out of 124 neurons, 8 mice) were CFC-R while 45% of PV interneurons were CFC-NR (56 out of 124 neurons, 8 mice; Figure 1F). However, in 5XFAD/PV-Cre mice, only 18% of PV interneurons (19 out of 104 neurons, 7 mice) were CFC-R while 82% of PV interneurons were CFC-NR (85 out of 104 neurons, 7 mice; Figure 1F). Overall, the population of CFC-R PV interneurons that responded to CFC in each mouse was significantly lower in 5XFAD/PV-Cre mice compared to PV-Cre mice (Chi squared text, *p* < 0.001; Figure 1F). This lower CFC-R ratio was correlated with the freezing ratio during CFM retrieval session (Figure 1G). These results indicate that encoding of CFM by RSC PV interneuron is disrupted in 5XFAD/PV-Cre mice.

**Figure 1.**
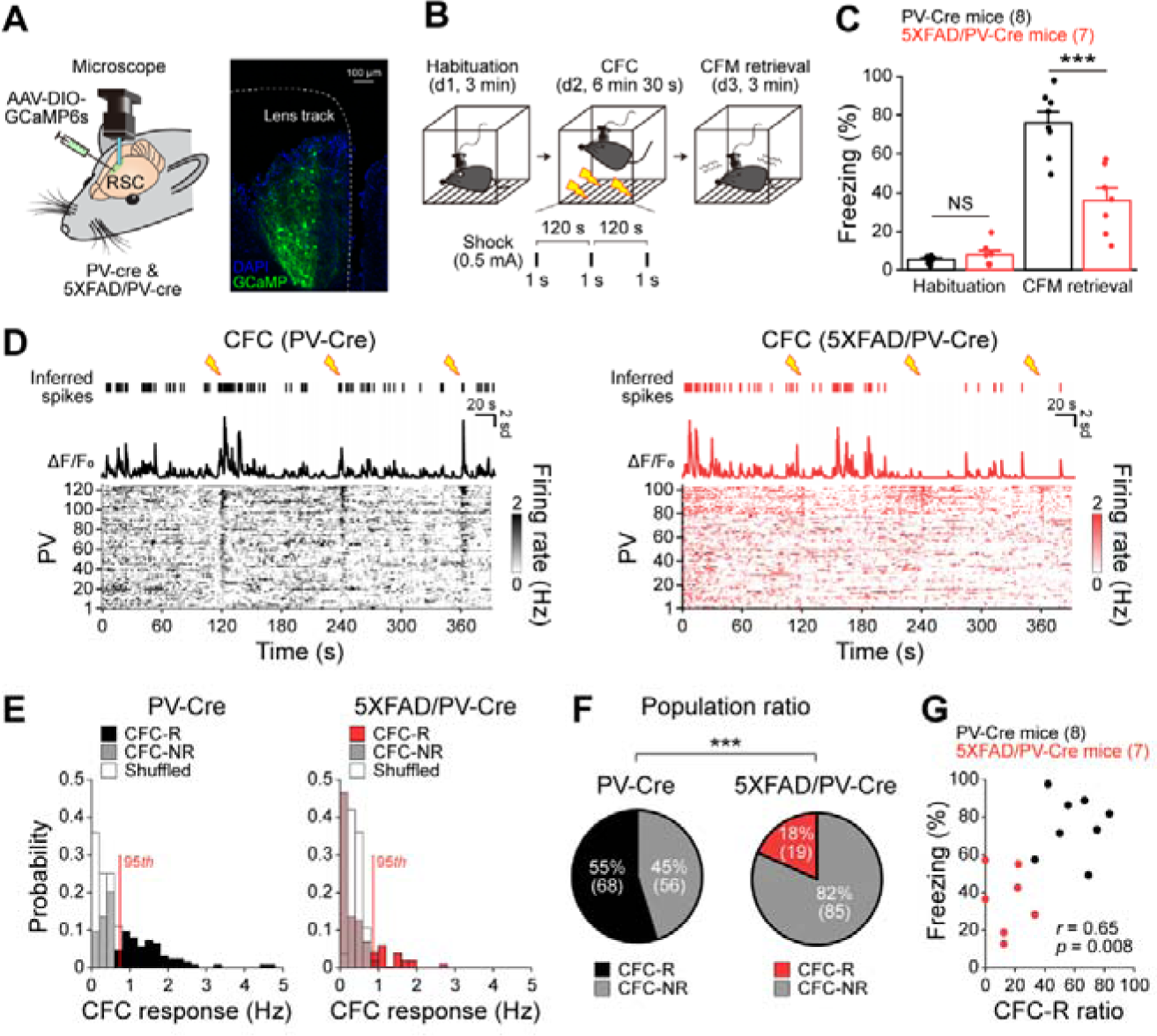
Encoding of CFM by PV interneurons in the granular retrosplenial cortex is impaired in 5XFAD/PV-Cre mice. (**A**) Experimental design for Ca^2+^ imaging from mice using a miniaturized microscope during contextual fear memory (CFM) task. AAV-DIO-GCaMP6s virus was injected in the granular retrosplenial cortex (RSC) of PV-Cre and 5XFAD/PV-Cre mice followed by a Gradient-index (GRIN) lens implant (left). Fluorescent image (right) of virally expressed GCaMP6s on the granular RSC PV interneurons. (**B**) Schematic illustration of CFM task. (**C**) Averaged freezing behavior percentage of PV-Cre mice and 5XFAD/PV-Cre mice during habituation and CFM retrieval sessions of CFM task. (**D**) Representative raw Ca^2+^ signals (△F/F_0_) of a single PV interneuron in RSC of PC-Cre mouse (black) or 5XFAD/PV-Cre mouse (red) from which deconvoluted spikes were extracted during CFC session (Inferred spike). RSC neuronal firing rate map of total imaged PV interneuron during CFC session. (**E**) Probability distributions of spike firing rate of CFC-responsive (CFC-R) and CFC non-responsive (CFC-NR) PV interneurons from PV-Cre mice or 5XFAD/PV-Cre mice in response to electric foot shocks (CFC response) plotted together with the randomly shuffled CFC response for comparison (white). Red line: 95th percentile of randomly shuffled CFC response. (**F**) Pie chart showing population ratio of CFC-R and CFC-NR PV interneurons. (**G**) CFC-R PV interneuron ratio versus averaged freezing behavior percentage of PV-Cre mice and 5XFAD/PV-Cre mice. Data are represented as mean ± SEM. \*\*\**p* < 0.001, n.s.: *p* > 0.05 for unpaired Student’s *t* test (C). \*\*\**p* < 0.001 for Chi-squared test of proportions (F).

### Selective upregulation of CFM-encoding PV interneuron ensemble dynamics during CFM retrieval is disrupted in 5XFAD/PV-Cre mice

We next investigated how the recruitment of CFC-R PV interneurons shapes the inhibitory ensemble dynamics underlying CFM retrieval in PV-Cre control mice, and consequently the lower number of CFC-R PV interneurons in 5XFAD/PV-Cre mice affects inhibitory ensemble dynamics. Analysis of PV interneuron ensemble activity during CFM retrieval in PV-Cre mice (Figure 2A) revealed that only CFC-R PV interneurons, but not CFC-NR PV interneurons, selectively increased their spike firing rate during the CFM retrieval session compared to the habituation session (Figure 2B). In contrast, in 5XFAD/PV-Cre mice, no changes were observed in spike firing rates during CFM retrieval and habituation sessions in neither CFC-R nor CFC-NR PV interneurons (Figure 2A, B). Thus, the selective increase in CFC-R PV interneuron ensemble activity during CFM retrieval session observed in PV-Cre mice was completely absent in 5XFAD/PV-Cre mice.

**Figure 2.**
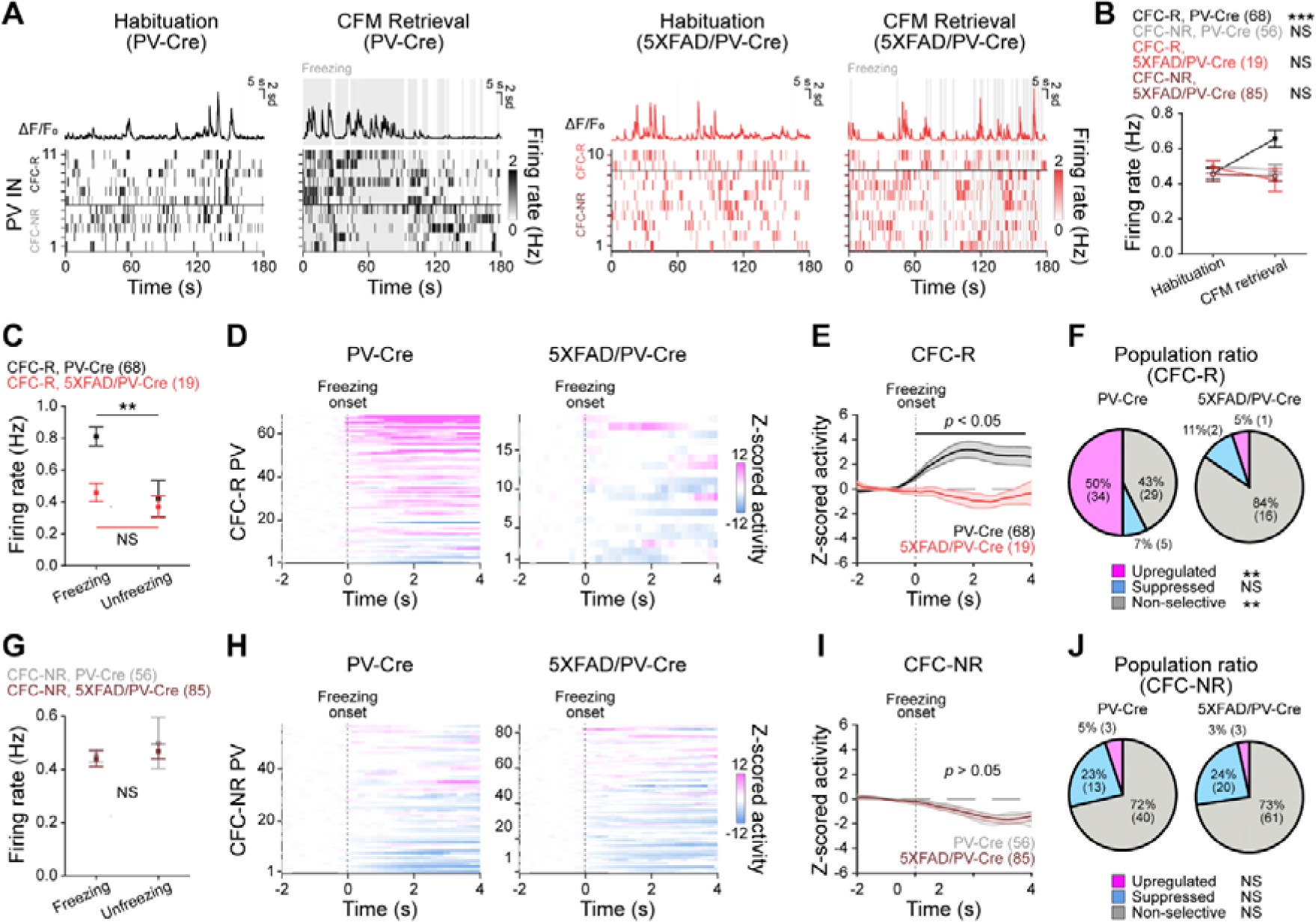
Selective upregulation of CFM-encoding PV interneuron ensemble dynamics in the granular retrosplenial cortex during CFM retrieval is disrupted in 5XFAD/PV-Cre mice. (**A**) Representative raw Ca^2+^ signals (△F/F_0_) of a single PV interneuron in the granular retrosplenial cortex (RSC) of PC-Cre mouse (black) or 5XFAD/PV-Cre mouse (red) during habituation and CFM retrieval sessions. RSC neuronal firing rate map of representative PV interneurons from the same mouse for each session. (**B**) Averaged firing rate of CFC responsive (CFC-R) and CFC non-responsive (CFC-NR) PV interneurons for each session. (**C**) Averaged firing rate of CFC-R PV interneurons during freezing and unfreezing epochs of CFC retrieval session. (**D**) Z-scored neuronal activity map of CFC-R PV interneurons at the onset of freezing epoch. (**E**) Averaged Z-cored neuronal activities of CFC-R PV interneuron at the onset of freezing epoch. Solid line: mean Z-scored activity. Shade: SEM. Black horizontal line: significant differences according to the Permutation test. (**F**) Population ratio of upregulated, suppressed, and non-selective CFC-R PV interneuron according to the mean Z-score at the onset of freezing epoch. (**G**-**J**) Same with (C-F), but for CFC-NR interneurons from PV-Cre mice or 5XFAD/PV-Cre mice. Data are represented as mean ± SEM. \*\*\**p* < 0.001, \*\**p* < 0.01, n.s.: *p* > 0.05 for paired Student’s *t* test (B, C, G). Permutation test (E, I). \*\**p* < 0.01, n.s.: *p* > 0.05 for Chi-squared test of proportions (F, J).

As freezing behavior is considered to be the behavioral correlate of CFM retrieval, we further investigated how ensemble activity of PV interneurons is related to freezing and unfreezing epochs during the CFM retrieval session. CFC-R PV interneuron selectively increased their spike firing rate during the freezing epoch in PV-Cre mice compared to the unfreezing epoch (Figure 2C, black), while in 5XFAD/PV-Cre mice, CFC-R PV interneurons spike firing rates were not different in freezing and non-freezing epochs (Figure 2C, red). To further characterize the ensemble dynamics of CFC-R PV interneurons during the freezing and unfreezing epochs, Z-scored neuronal activities were analyzed. We found that the majority of CFC-R PV interneurons (50%, 34 out of 68 neurons; Figure 2D-F, Figure S2) in PV-Cre mice showed upregulated Z-scored neuronal activities during the freezing epochs while subpopulations of CFC-R PV interneurons showed either no changes (43%, 29 out of 68 neurons; Figure 2D, Figure S2) or suppressed Z-scored neuronal activities (7%, 5 out of 68 neurons; Figure 2D. Figure S2). Subpopulations of CFC-R PV interneurons that were upregulated during the freezing epoch of CFM retrieval were selectively suppressed in Z-scores during the unfreezing epoch of CFM retrieval, showing bi-directional modulation of ensemble activity depending on the behavioral states (Figure S3). However, such freezing/unfreezing-selective CFC-R PV interneurons responses were independent of locomotion *per se*, as they showed low correlation between their neural activities and mouse velocity (Figure S4). In contrast, in 5XFAD/PV-Cre mice, the majority of CFC-R PV interneurons in 5XFAD/PV-Cre mice showed no changes in Z-scored neuronal activities during both the freezing (84%, 16 out of 19 neurons; Figure 2D-F, Figure S2) and unfreezing epochs (84%, 16 out of 19 neurons; Figure S2, S3). Also, analysis of CFC-NR PV interneurons showed no differences in spike firing rate (Figure 2G) nor Z-scored neural activities during freezing epochs of CFM retrieval session in PV-Cre mice and in 5XFAD/PV-Cre mice (PV-Cre: 72%, 40 out of 56 neurons, 5XFAD/PV-Cre: 73%, 61 out of 85 neurons; Figure 2H-J, Figure S2). Together, these results suggest that selective upregulation of CFC-R PV interneuron ensemble dynamics during the freezing epoch plays a key role in CFM retrieval in PV-Cre control mice while disruption of CFC-R PV interneuron ensemble dynamics in 5XFAD/PV-Cre mice may explain deficits in their CFM retrieval.

### CFM-induced synaptic plasticity of PV interneuron-mediated inhibition is impaired in 5XFAD/PV-Cre mice

As CFC-R PV interneuron ensemble activity was selectively and dynamically upregulated during CFM retrieval session in PV-Cre mice, it is likely that the CFC-induced RSC PV interneuron ensemble activity was mediated by the synaptic strengthening of PV interneuron-mediated inhibition in the network. To directly test this hypothesis, we performed whole-cell voltage-clamp recordings to measure the changes in PV interneuron-mediated inhibition following CFC *in vitro*. Mice were injected with AAV-ChR2-DIO-eYFP and AAV-CaMKII-mCherry in the granular RSC (Figure 3A, top), and from brain slices cut from either naive mice (Naive) or mice that underwent CFC (post-CFC), PV interneuron-evoked inhibitory postsynaptic currents (PV-eIPSCs) were recorded from mCherry-expressing excitatory neurons *in vitro* by stimulating ChR2-expressing PV interneurons using blue light (470 nm, Figure 3A, bottom). In response to incrementing amplitudes of single-pulse blue light stimulation, PV-eIPSCs in the stimulus-response (S-R) curve revealed that PV-eIPSCs amplitudes in the naive 5XFAD/PV-Cre mice was significantly decreased compared to those in the naive PV-Cre mice, indicating the reduction of responsiveness in PV interneurons in 5XFAD/PV-Cre mice even in naive condition (Figure 3B, C). PV-eIPSCs amplitudes were significantly increased in brain slices cut from PV-Cre mice post-CFC compared to those from the naive PV-Cre mice (Figure 3C, black), but such changes were absent in 5XFAD/PV-Cre mice (Figure 3C, red). These results indicate that not only PV interneuron responsiveness is impaired in 5XFAD/PV-Cre mice in the naive condition but also that potentiation of PV interneuron-mediated inhibition at the PV-to-excitatory neuron synapse is impaired. To further investigate the short-term synaptic changes of PV interneuron-mediated inhibition, paired-pulse ratio (PPR) was measured by delivering two pulses of blue light stimulation at 20 Hz to ChR2-expressing PV interneurons (Figure 3D, E). Paired-pulse facilitation of PV-eIPSCs in the naive PV-Cre mice was converted to paired-pulse depression in PV-Cre mice post-CFC (Figure 3D, E, black), indicating that CFC induced presynaptic strengthening of PV-eIPSCs to excitatory neurons. However, paired-pulse facilitation in the naive 5XFAD/PV-Cre mice was unaltered in 5XFAD/PV-Cre mice even post-CFC (Figure 3D, E, red). Based on the S-R curve and PPR analyses, it can be inferred that CFC-induced enhancement of PV interneuron responsiveness and synaptic strengthening of PV interneuron-mediated inhibition may have contributed to the upregulation of PV interneurons ensemble dynamics during the freezing epoch of CFM retrieval. In contrast, the reduced responsiveness and synaptic dysfunctions of PV interneurons may have disrupted the recruitment of CFC-R PV interneurons during CFM encoding, which leads to the failure in shaping the ensemble dynamics underlying CFM retrieval in 5XFAD/PV-Cre mice.

**Figure 3.**
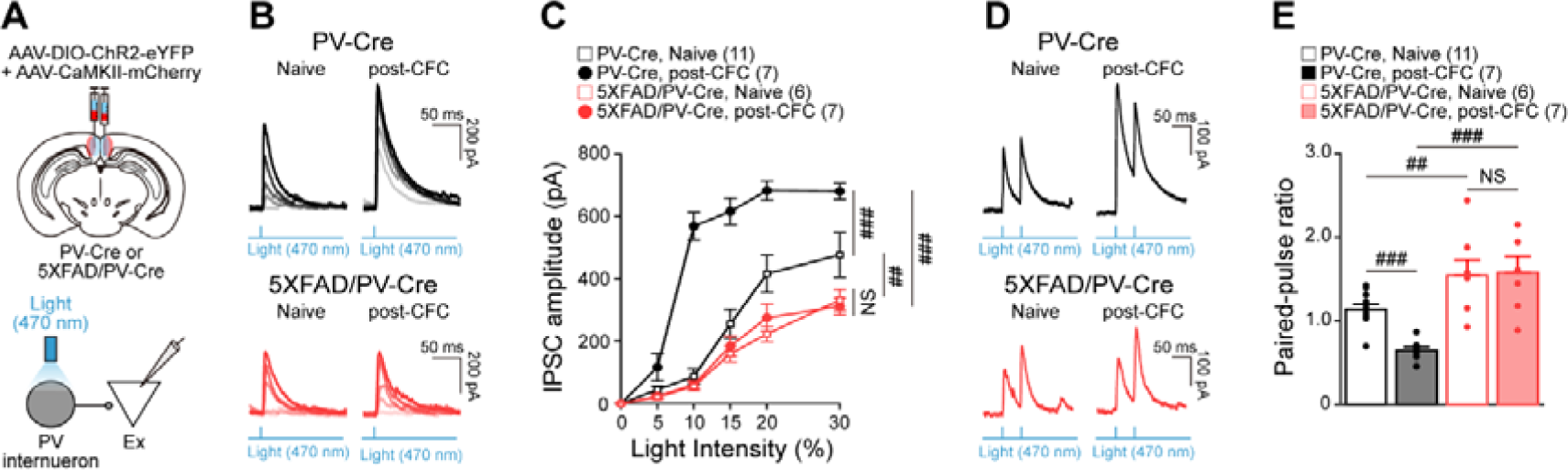
CFM-induced potentiation of PV interneuron-mediated inhibition in the granular retrosplenial cortex is impaired in 5XFAD/PV-Cre mice. (**A**) Experimental design (top) for simultaneous virus injection of AAV-DIO-ChR2-eYFP and AAV-CaMKII-mCherry in the granular retrosplenial cortex (RSC). Schematic showing *in vitro* whole-cell voltage-clamp recordings from excitatory neurons (Ex) during blue light (470 nm) stimulation of ChR2-expressing PV interneurons in RSC. (**B**, **C**) Representative PV interneuron-evoked inhibitory postsynaptic currents (IPSCs) in response to different light stimulation powers (B) and stimulus-response curve (C), recorded from RSC Ex of PV-Cre (top) and 5XFAD/PV-Cre mice (bottom) without (Naive) or after CFC (post-CFC). (**D**, **E**) Representative PV interneuron-evoked inhibitory postsynaptic currents (IPSCs) in response to light stimulations (2 pulses, 20 Hz) (D) and paired-pulse ratio of the 2nd IPSC/1st IPSC (E), recorded from RSC Ex for each condition. Data are represented as mean ± SEM. ###*p* < 0.001, ##*p* < 0.01, n.s.: *p* > 0.05 for two-way (C) and one-way ANOVA with *post hoc* Tukey’s test (E).

### Optogenetic activation of PV interneurons during CFM-encoding restores CFM retrieval in 5XFAD/PV-Cre mice

So far, we found that failure of PV interneurons in 5XFAD/PV-Cre mice to encode CFM led to the disruption of ensemble dynamics required for CFM retrieval. Thus, we next tested whether disruption of CFM encoding in PV interneurons in PV-Cre mice could mimic the CFM deficits observed in 5XFAD/PV-Cre mice. We injected AAV-DIO-NpHR3.0-eYFP to optogenetically silence NpHR3.0-expressing PV interneurons using yellow light (595 nm) delivered through optic fibers implanted in the granular RSC (Figure 4A, top) when electric foot shocks were delivered during CFC (Figure 4B). In these mice, the freezing ratio during CFM retrieval session significantly decreased compared to that in PV-Cre mice, which was similar to the freezing ratio observed in 5XFAD/PV-Cre mice (Figure 4C). These results indicate that silencing of PV interneurons to disrupt the recruitment of CFC-R PV interneurons during CFM encoding could mimic the deficits in CFM retrieval observed in 5XFAD/PV-Cre mice. If disruption of CFM encoding by PV interneurons can explain deficits in CFM retrieval, it is likely that increased recruitment of PV interneurons during CFM encoding in 5XFAD/PV-Cre mice could restore CFM retrieval. To test this hypothesis, in 5XFAD/PV-Cre mice, we injected AAV-Syn-Flex-Chrimson-tdTomato to granular RSC and activated Chrimson-expressing PV interneurons using yellow light through optic fibers implanted in the RSC (Figure 4A, bottom) when electric foot shocks were delivered during CFC (Figure 4B). Interestingly, yellow light stimulation of Chrimson-expressing PV interneurons increased the freezing ratio during CFM retrieval session in 5XFAD/PV-Cre mice to the similar level as that in PV-Cre mice (Figure 4C). Overall, our optogenetic manipulation results reveal that the recruitment of CFM-encoding RSC PV interneurons during CFC can gate CFM retrieval in the healthy brain and optogenetic activation of dysfunctional RSC PV interneurons during CFM encoding in 5XFAD/PV-Cre mice is sufficient to restore memory deficits.

**Figure 4.**
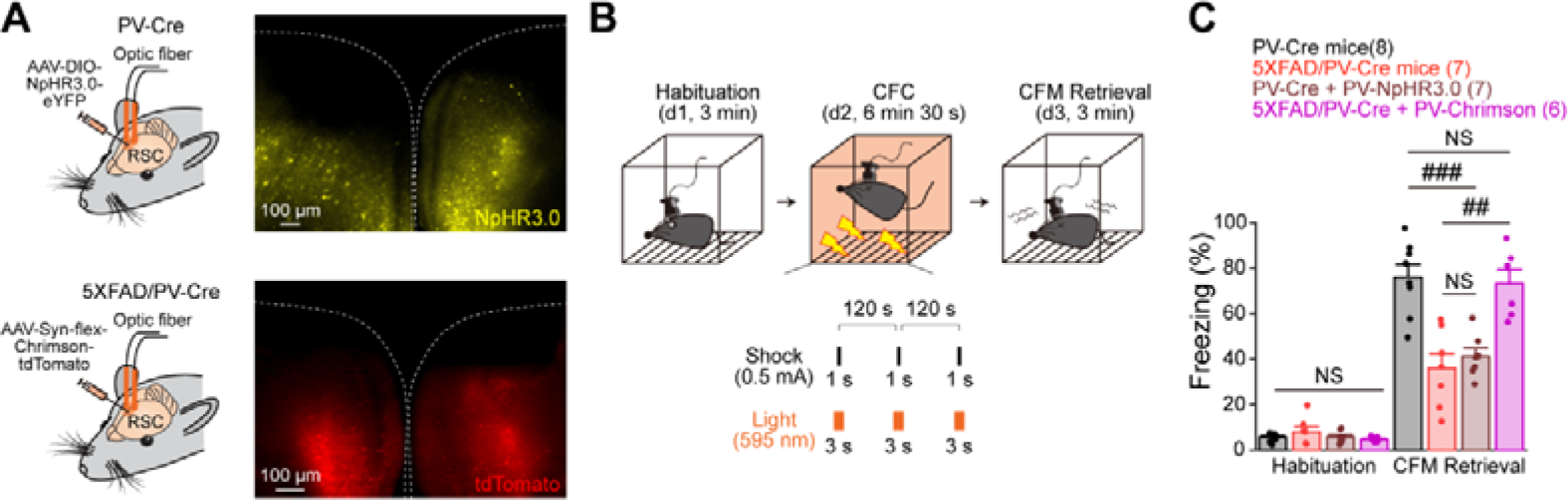
Optogenetic activation of PV interneuron in the granular retrosplenial cortex during CFM encoding restores CFM retrieval in 5XFAD/PV-Cre mice. (**A**) Experimental design for optogenetic inactivation (top, left) or activation (bottom, left) of PV interneurons in the granular retrosplenial cortex (RSC) during CFC. AAV-DIO-NpHR3.0-eYFP (top) or AAV-Syn-flex-Chrimson-tdTomato (bottom) was injected in RSC of PV-Cre mice or 5XFAD/PV-Cre mice, respectively, followed by optic fiber implant. Fluorescent images of virally expressed NpHR3.0 (top, right) or Chrimson (bottom, right) on the granular RSC PV interneurons. (**B**) Schematic illustration of CFM task with optogenetic inactivation or activation of RSC PV interneuron during CFC. (**C**) Averaged freezing behavior percentage of each condition during habituation and CFM retrieval sessions of CFM task. Data are represented as mean ± SEM. ###*p* < 0.001, n.s.: *p* > 0.05 for one-way ANOVA with post hoc Tukey’s test (C).

### Concluding Remarks

By combining *in vivo* and *in vitro* neural circuit analysis, we found that RSC PV interneuron can encode CFM (Figure 1) and CFM-encoding PV interneuron ensemble dynamics were selectively upregulated during the CFM retrieval session (Figure 2), which was manifested by the potentiation of PV interneuron-mediated inhibition at the synaptic level after CFC (Figure 3). However, in 5XFAD/PV-Cre mice, synaptic dysfunctions of RSC PV interneurons disrupted the encoding of CFM in RSC PV interneurons, which consequently led to the failure in shaping the ensemble dynamics underlying CFM retrieval in 5XFAD/PV-Cre mice (Figure 1-3). Optogenetic activation of PV interneurons in 5XFAD/PV-Cre during CFC to recruit more CFM-encoding PV interneurons could fully restore deficits in CFM retrieval in 5XFAD/PV-Cre mice while optogenetic silencing of PV interneurons in PV-Cre mice to interfere with the recruitment of CFM-encoding PV interneurons could mimic the deficits in CFM retrieval in 5XFAD/PV-Cre mice (Figure 4). Together, these results suggest that the ensemble dynamics of CFM-encoding RSC PV interneuron represents CFM retrieval and that dysfunction of RSC PV interneurons disrupts ensemble dynamics underlying CFM encoding and retrieval, leading to memory deficits in 5XFAD mouse model of AD.

Our results demonstrate that RSC PV interneurons can represent encoding and retrieval of memory, which is in line with the many previous studies showing that GABAergic interneurons play a key role in encoding memory^8–12^. Moreover, our results suggest that RSC PV interneurons also display properties of inhibitory memory engram^9^ as 1) they were activated during CFM encoding, (Figure 1), 2) CFM-encoding RSC PV interneurons were selectively re-activated during CFM retrieval (Figure 2), and 3) synaptic strengthening of PV interneuron-mediated inhibition post-CFC was observed (Figure 3). Thus, it is possible that RSC PV inhibitory memory engrams identified here could perform many of the suggested functions of inhibitory engrams such as fine-tuning of excitatory engrams in RSC^14, 15^ by controlling their size^11, 24^, suppressing the irrelevant excitatory neurons^9^, and activating the appropriate excitatory memory engrams through disinhibition^8, 10, 12, 25, 26^. As inhibitory engrams are suggested to be “a negative image” of excitatory engrams, it can be predicted that the ensemble dynamics of RSC excitatory neurons will be downregulated during freezing epoch while upregulated during the unfreezing epoch of CFM retrieval. In fact, subpopulations of RSC excitatory neurons are known to process spatial context information during animal’s navigating movement^13^, especially with the recent discovery of egocentric boundary vector/border cells in the RSC^27, 28^. Thus, spatial context information processed by RSC excitatory neurons may be activated during the unfreezing-epoch during CFM-retrieval, which in turn may activate CFC-R PV interneurons to initiate freezing behavior. Also, as RSC is known to project to motor cortex^29^, PV interneurons may inhibit RSC projection to motor cortex to initiate freezing activity, which will be interesting to explore in the future.

In our study, optogenetic activation of RSC PV interneuron to recruit more CFC-R PV interneurons in 5XFAD/PV-Cre mice restored CFM retrieval, further underscoring the critical roles of CFM-encoding RSC PV interneurons for successful CFM retrieval (Figure 4). Such result may explain how dysfunction of PV interneurons has been associated with the pathological features of AD^1–3, 30–37^ and how manipulations of PV interneurons in AD through optogenetic stimulation^1, 2, 30, 31^, genetic manipulation^33^, and PV interneuron transplant^34^ ameliorated memory deficits in AD. In addition to the disruption of inhibitory ensemble dynamics, dysfunction of RSC PV interneurons may have led to excitation-inhibition (E-I) imbalance^4, 5, 38–40^ and the hyperexcitability^32, 33, 41–43^ of neural network in AD, which have been reported to disrupt the neural correlates of memory such as the network oscillation^1, 2, 30–33, 40, 44–49^ and synaptic plasticity^2, 39, 50–54^ in animal models of AD or in AD patients. Thus, elucidating how impairments of network oscillation and synaptic plasticity are associated with disruption of inhibitory engrams may further our understanding on the dysfunctions of neural correlates of memory in AD.

Overall, our results provide experimental evidence for how CFM encoding in RSC PV interneuron contributes to shaping the ensemble dynamics underling CFM retrieval in the cortical network and how dysfunction of RSC PV interneurons disturbs the neural representation of CFM encoding and retrieval, leading to memory deficits in AD. Our results provide insight into the neural circuit mechanisms of memory processing and point towards potential therapeutic targets in AD.

## STAR METHODS

- KEY RESOURCES TABLE
- RESOURCE AVAILABILITY
  ✓ Lead contact
  ✓ Materials availability
  ✓ Data and code availability
- EXPERIMENTAL MODEL AND SUBJECT DETAILS
  ✓ Animals
- METHODS DETAILS
  ✓ Virus and stereotaxic surgery for *in vivo* Ca^2+^ imaging
  ✓ Virus and stereotaxic surgery for *in vitro* optogenetic experiments
  ✓ Virus and stereotaxic surgery for *in vivo* optogenetic experiments
  ✓ *In vivo* Ca^2+^ imaging and signal processing
  ✓ Contextual fear memory task
  ✓ Histology and immunohistochemistry
  ✓ Neural selectivity analysis
  ✓ *In vitro* RSC slice preparation
  ✓ *In vitro* whole-cell patch-clamp recordings
- QUANTIFICATION AND STATISTICAL ANALYSIS
  ✓ Statistics

## SUPPLEMENTARY INFORMATION

Figure S1 – S4

## ACKNOWLEDGMENTS

“This research was supported by a grant of the Korea Dementia Research Project through the Korea Dementia Research Center (KDRC), funded by the Ministry of Health & Welfare and Ministry of Science and ICT, Republic of Korea (grant number: HU20C0233).” We thank Mr. Yoonsoo Yeo for help with animal behavior experiments.

## AUTHOR CONTRIBUTIONS

K.P., M.M.K., and J.K. conceived of the experiment. K.P. collected and analyzed the data.

J.K. wrote the first draft of the manuscript. K.P., M.M.K. and J.K. edited the manuscript.

## INCLUSION AND DIVERSITY

We support inclusive, diverse, and equitable conduct of research

## STAR METHODS

### KEY RESOURCES TABLE

| REAGENT or RESOURCE | SOURCE | IDENTIFIER |
| --- | --- | --- |
| **Experimental models: Organisms/strains** |  |  |
| PV-Cre mice | Jackson Laboratory | https://www.jax.org/strain/017320 |
| 5XFAD mice | Jackson Laboratory | https://www.jax.org/strain/006554 |
| 5XFAD/PV-Cre mice | This paper |  |
| **Bacterial and virus strains** |  |  |
| AAV1-EF1a-DIO-GCaMP6s-P2A-nls-dTomato | Addgene | https://www.addgene.org/51082/ |
| AAV9-Ef1a-DIO-ChR2(E123T/T159C)-eYFP | Addgene | https://www.addgene.org/35509/ |
| AAV5-CaMKII-mCherry | Addgene | https://www.addgene.org/114469/ |
| AAV5-Ef1a-DIO-NpHR3.0-eYFP | Addgene | https://www.addgene.org/26966/ |
| AAV5-Syn-flex-rc[ChrimsonR-tdTomato] | Addgene | https://www.addgene.org/62723/ |
| **Antibodies** |  |  |
| Rabbit anti-beta amyloid 1–42 | Abcam | https://www.abcam.com/products/primary-antibodies/beta-amyloid-1-42-antibody-moc64-ab201060.html |
| Donkey anti-rabbit | Jackson ImmunoResearch | https://www.jacksonimmuno.com/catalog/products/711-167-003 |
| **Software and algorithms** |  |  |
| MATLAB R2021a | Mathwork | https://kr.mathworks.com/ |
| Inscopix Data Processing Software | Inscopix | https://iq.inscopix.com/software |
| Ethovision XT 16 | Noldus | https://www.noldus.com/ethovision-xt |
| Adobe Illustrator 2021 | Adobe | https://www.adobe.com/ |
| Igor Pro | WaveMetrics | https://www.wavemetrics.com/downloads/current |
| **Other** |  |  |
| Miniaturized microscope | Inscopix | https://www.inscopix.com/nvoke |
| GRIN lens | Inscopix | https://www.inscopix.com/lenses-viruses |
| Optic cannula | RWD | https://www.rwdstco.com/product-item/fiber-optic-cannulae-with-ceramic-ferrule/ |
| LED driver | Thorlabs | https://www.thorlabs.com/newgrouppage9.cfm?objectgroup_id=2616 |

### RESOURCE AVAILABILITY

#### Lead Contract

Request for further information and resources should be directed to and will be fulfilled by the lead contract Dr. Jeehyun Kwak.

#### Materials availability

This study did not generate unique reagents.

#### Data and code availability

All data reported in this paper and the code employed for the analysis will be shared by the lead contract upon request. Any additional information required to reanalyze the data reported in this paper is available from the lead contact upon request.

## EXPERIMENTAL MODEL AND SUBJECT DETAILS

### Animals

Total two different lines of male mice (6 – 9 months old) including the control mice (PV-Cre mice, #017320, Jackson Laboratory, USA) and Alzheimer’s disease (AD) mice model (5XFAD/PV-Cre mice^31^) were used with the approval of the Institutional Animal Care and Use Committee (IACUC) at Korea University (KUIACUC-2020-0099, KUIACUC-2021-0052). 5XFAD/PV-Cre mice were generated by crossbreeding of PV-Cre mice and 5XFAD mice^55^ (#034840, Jackson Laboratory, USA). All mice were individually housed in a temperature- and humidity-controlled vivarium, which was kept in a 12 h/12 h light/dark cycle. All experiments were performed during the dark cycle. Food and water were available ad libitum.

## METHODS DETAILS

### Virus and stereotaxic surgery for *in vivo* Ca^2+^ imaging

For surgical procedure, mice were deeply anesthetized using 2% isoflurane (2 ml/min flow rate) and head-fixed into a stereotaxic frame (51730D, Stoelting Co., USA). For *in vivo* Ca^2+^ imaging of PV interneurons from PV-Cre or 5XFAD/PV-Cre mice (Figure 1, 2, Figure S2-4), AAV1-EF1a-DIO-GCaMP6s-P2A-nls-dTomato (#51082, Addgene, USA) in solution (5 x 10^12^ virus molecules/ml diluted with saline at 1:1 ratio) was used. Virus was injected into three sites (500 nl per injection site) along the dorsoventral axis of the retrosplenial cortex (RSC) (AP: –2.5 mm, ML: –0.3 mm, DV: –0.8, –0.6, and –0.4 mm) using a Hamilton syringe (#87930, Hamilton, USA) controlled by an automated stereotaxic injector (100 nl/min; #53311, Stoelting Quintessential Injector, Stoelting Co., USA). The syringe was left at the injection coordinates for more than 5 min to allow viral diffusion.

After viral injection, a Gradient-index (GRIN) lens (1 mm diameter, 4 mm length; #1050-004605, Inscopix, USA) was implanted to RSC (AP: –2.5 mm; ML: –0.3 mm; DV: –0.5 mm) and fixed to the skull with dental cement (Self Curing, Vertex, Netherlands). After two weeks of recovery time following the surgery, mice were deeply anesthetized using 2% isoflurane (2 ml/min flow rate) to attach a magnetic baseplate (#1050-004638, Inscopix, USA), on top of which a miniaturized microscope (nVoke, Inscopix, USA) was mounted. If GCaMP6s-expressing neurons were visible in the field of view (FOV), baseplate was fixed to the skull using dental adhesive resin cement (Super bond, Sun Medical, Japan) to perform *in vivo* longitudinal Ca^2+^ imaging of the same FOV. After baseplating, at least one week of recovery time was allowed before performing *in vivo* Ca^2+^ imaging.

### Virus and stereotaxic surgery for *in vitro* optogenetic experiments

For *in vitro* optogenetic stimulation of PV interneurons from PV-Cre or 5XFAD/PV-Cre mice (Figure 3), both AAV9-Ef1a-DIO-ChR2(E123T/T159C)-eYFP (#35509, Addgene, USA) in solution (1 x 10^13^ virus molecules/ml diluted with saline at 3:1 ratio) and AAV5-CaMKII-mCherry (#114469, Addgene, USA) in solution (7 x 10^12^ virus molecules/ml diluted with saline at 3:1 ratio) were used following the same virus injection protocols for *in vivo* Ca^2+^ imaging. After viral injection, at least two week of recovery time was allowed before performing *in vitro* optogenetic experiments.

### Virus and stereotaxic surgery for *in vivo* optogenetic experiments

For *in vivo* optogenetic inactivation/activation of PV interneurons from PV-Cre or 5XFAD/PV-Cre mice (Figure 4), AAV5-Ef1a-DIO-NpHR3.0-eYFP (#26966, Addgene, USA) in solution (1 x 10^13^ virus molecules/ml diluted with saline at 3:1 ratio) or AAV5-Syn-flex-rc[ChrimsonR-tdTomato] (#62723, Addgene, USA) in solution (5 x 10^12^ virus molecules/ml diluted with saline at 3:1 ratio) were used, respectively, following the same virus injection protocols for *in vivo* Ca^2+^ imaging with slight modification of injection site (AP: ±2.5 mm, ML: –0.3 mm, DV: –0.8, –0.6, and –0.4 mm). After viral injection, two optic cannulas (400 μm core diameter, 2 mm length; #087-00017-00, RWD, China) was implanted to both right and left RSC (AP: ±2.5 mm; ML: –0.3 mm; DV: –0.5 mm) and fixed to the skull with dental cement (Self Curing, Vertex, Netherlands). After optic fiber implant, at least two week of recovery time was allowed before performing *in vivo* optogenetic experiments.

### *In vivo* Ca^2+^ imaging and signal processing

Prior to *in vivo* Ca^2+^ imaging during behavior task, mice were adapted to the head-mounted miniaturized microscope connected to the commutator systems (Inscopix, USA) for at least five days. All experiments were performed under the luminescence of 30 Lux unless it is mentioned otherwise.

*In vivo* Ca^2+^ imaging data from GCaMP6s-expressig RSC neurons were acquired using nVoke acquisition software (Inscopix, USA) at a sampling rate of 20 Hz (exposure time: 50 ms). An optimal LED power (0.1 – 0.7 mW/mm^2^) and digital focus (0 – 1000) was selected for each mouse based on GCaMP6s expression on neurons in the FOV and the same parameters were used for each mouse for longitudinal imaging.

Ca^2+^ image processing was performed using Inscopix Data Processing Software (Inscopix, USA), following other studies^56, 57^. For all data, motion correction was applied using an image registration method^56, 58^. Mean fluorescence signal value (F) of each pixel across the entire Ca^2+^ image video was used to calculate the reference, termed F_0_. Then, changes in fluorescence signal value (△F) of each pixel were normalized by F_0_ to acquire △F/F_0_. Putative neurons and their Ca^2+^ signals were isolated using automated cell-segmentation algorithm on △F/F_0_ data, which is based on principal component analysis-independent component analysis^56, 59^.

Putative spikes were inferred from △F/F_0_ signal using spike deconvolution algorithm called “Online Active Set methods to Infer Spikes”^57, 60–62^. Spikes with an amplitude less than 3 signals-to-noise ratio of deconvolved Ca^2+^ signals were removed from the data.

### Contextual fear memory task

Contextual fear memory (CFM) task^23^ was performed in a rectangular shaped conditioning chamber (width: 30 cm, length: 26 cm, height: 33 cm, Coulbourn Instruments, USA) with stainless-steel grid floor composed of 16 grid bars. The grid floor was connected to a precision animal shocker (Coulbourn Instruments, USA) set to deliver electrical foot shocks (0.5 mA, 1 s-long). On the first day of CFM task (habituation), mice were habituated in the conditioning chamber for 3 min. On the second day, mice were placed in the same chamber for contextual fear conditioning (CFC), consisting of 120 s pre-stimulus period followed by delivery of three aversive electric foot shocks (120 s inter-stimulus intervals) to form associative memory between the context of the chamber and the electric foot shocks. On the last day, mice were returned to the same chamber for 3 min without the delivery of electric foot shocks to assess CFM retrieval, measured as freezing behavior. Freezing behavior was defined based on the immobility of mice which was captured by detecting the time epochs showing the video pixels of a detected mouse between the behavior data sample points less than threshold (0.5 – 3%)^63^. The threshold of video pixel change for each mouse was decided based on the visual inspection.

For *in vivo* optogenetic stimulation of PV interneurons during CFM task (Figure 4), optic cannulas were connected to LED driver (T-Cube^TM^ LED driver, Thorlabs, USA) via patch cables connected to a FC/PC rotary joint (RJ1, Thorlabs, USA). Mice were adapted to the attached patch cables at least five days before CFM task. During whole CFM task, patch cables were attached and 595 nm yellow light stimulations (3 s-long) were delivered 1 s preceding each electrical foot shock to optogenetically stimulate PV interneurons during CFC. Light intensity was set to output 5 mW at the tip of the optic fibers.

After each session of CFM task, the conditioning chamber was cleaned using 70% ethanol to eliminate any odors. Mice behavior was recorded by a video camera placed on the ceiling above the chambers or arenas at a sampling rate of 7.5 Hz. All behavior data including video pixel changes and velocity were analyzed using Ethovision XT 16 program (Noldus, Netherlands).

### Histology and immunohistochemistry

To confirm the position of GRIN lens track or optic fiber track and viral expression in granular RSC PV interneurons after Ca^2+^ imaging or *in vivo* optogenetic experiments, the brain was removed from the mice that were deeply anesthetized using Avertin and perfused with 10 ml chilled 4% paraformaldehyde (PFA, 158127, Sigma-Aldrich, USA) transcardially. The brain was fixed overnight in 4% paraformaldehyde and transferred to 30% sucrose solution in phosphate-buffered saline (PBS) for cryoprotection for 24 – 48 h. Brains were frozen in Optimum Cutting Temperature (OCT) compounds (Tissue-Tek O.C.T. Compound, SAKURA, Japan) at –50°C prior to cryosectioning in 100 μm-thick coronal slices on a cryostat (YD-2235, Jinhua Yidi Medical Appliance Co., China). Slices were washed three times with PBS to wash out the OCT compounds and mounted on slide glasses with an antifade mounting medium with DAPI (Vectashield, Vector Laboratories, USA). Ca^2+^ imaging locations or *in vivo* optogenetic stimulation locations were identified based on the position of GRIN lens track or optic fiber track, respectively. Expression of virus on PV interneurons was verified by detecting the fluorescent signal using a confocal microscope (LSM-700, ZEISS, Germany) and fluorescent microscope (DM2500, Leica, Germany).

To verify the deposition of beta-amyloid (Aβ) on the RSC area of brain slice from 5XFAD/PV-Cre mice, PFA-fixed brain slices were incubated in peroxidase buffer (0.3% H_2_O_2_ in 0.1 M PBS) for 20 min. Non-targeted antigens were blocked by incubation in 6% bovine serum albumin and 0.3% Triton X-100 in 0.1 M PBS for > 24 h at 4 °C, after which slices were incubated with primary anti-Aβ antibody (1:300 dilution in 0.1 M PBS, at 4 °C for 4 days)(anti-Aβ 1–42 antibody [mOC64]-Conformation-Specifc, #ab201060, Abcam, UK) and secondary antibody (1:500 dilution in 0.1 M PBS)(CyTM3 AfniPure Donkey Anti-Rabbit IgG (H +L), Jackson ImmunoResearch, USA). Slices were washed three times with PBS and mounted on slide glasses with an antifade mounting medium. Deposition of Aβ was verified by detecting the fluorescent signal using a confocal microscope (LSM-700, ZEISS, Germany) and fluorescent microscope (DM2500, Leica, Germany).

### Neural selectivity analysis

Ca^2+^ imaging data from Inscopix and behavior data from Ethovision XT 16 were synchronized using Noldus-IO box (Mini USB-IO box, Noldus, Netherlands) system, which were then analyzed with custom-made Matlab (Mathworks, USA) code. To analyze responsiveness of RSC PV interneurons to CFC-inducing electric foot shocks^64, 65^ (Figure 1), their spike firing rates in response to electrical foot shocks (CFC response) were calculated. A neuron was defined as CFC-responsive if the CFC response exceeded the 95^th^ percentile of randomly shuffled CFC response distributions all imaged RSC neurons. Shuffled CFC response distribution was generated by circularly time-shifting the spike trains of neurons with randomly selected period along the mouse behavior data with 100 repetitions following the method used in previous studies^66^. This random time-shuffling preserved the temporal structures of both Ca^2+^ signals and behavior data while disrupted the temporal correlation between them. Random time-shuffling was applied for all randomly shuffled distribution for other metric.

To analyze ensemble dynamics of RSC PV interneurons during freezing and unfreezing epochs^66, 67^ (Figure 2, Figure S2, S3), their neuronal firing rates were Z-scored in the time windows of -2 s to 4 s at the onset of freezing or unfreezing epochs^66^. Time window of -2 to 0 s at the freezing or unfreezing epochs were used as baseline periods for Z-scored analysis. A neuron was defined as upregulated if the mean Z-score of the neuronal activity during freezing or unfreezing epochs exceeded the 95^th^ percentile of randomly shuffled Z-score distributions of all imaged RSC neurons, respectively. On the other hand, A neuron was defined as suppressed if the mean Z-score of the neuronal activity during freezing or unfreezing epochs fell below the 5^th^ percentile of randomly shuffled Z-score distributions of all imaged RSC neurons, respectively. Other neurons were defined as non-selective to freezing or unfreezing epochs.

### *In vitro* RSC slice preparation

Mice were deeply anesthetized using 1.25% Avertin solution (8 g of 2, 2, 2-Tribromoethanol and 5.1 ml of 2-methyl-2-butanol in 402.9 ml saline, Sigma Aldrich, USA) at a delivery rate of 0.2 ml/10 g body weight and perfused with ice-cold cutting solution (containing (in mM): 180 sucrose, 2.5 KCl, 1.25 NaH_2_PO_4_, 25 NaHCO_3_, 11 glucose, 2 MgSO_4_, and 1 CaCl_2_ at pH[7.2–7.4 oxygenated with 95% O_2_/5% CO_2_). Coronal RSC slices (300 μm) were cut using a vibratome (VT 1000[S, Leica Microsystems, Germany). Slices were allowed to recover for 30 min in a mixture of cutting solution and artificial cerebrospinal fluid (aCSF, containing (in mM): 126 NaCl, 3 KCl, 1.25 NaH_2_PO_4_, 2 MgSO_4_, 2 CaCl_2_, 25 NaHCO_3_, and 10 glucose at pH[7.2–7.4 bubbled with 95% O_2_/5% CO_2_) solution at 1:1 ratio, after which the slices were further incubated in aCSF for at least 1 h at 30– 32 °C before performing *in vitro* electrophysiological recordings.

### *In vitro* whole-cell patch-clamp recordings

Slices were moved to a recording chamber filled with aCSF (30–32 °C), and RSC area was identified under the guidance of differential interference contrast microscopy (BW51W, Olympus). To perform *in vitro* optogenetic light stimulation (Figure 3), 470 nm blue light (X-Cite, Excelitas Technologies, USA) were delivered through the objective (x 40) of the microscope (BX51W, Olympus, Japan) which covered the 550 μm diameter circle of the RSC area with the center of the illumination positioned at the patch clamp electrode. To record PV interneuron-evoked inhibitory postsynaptic currents from excitatory neurons, channelrhodopsin-2 (ChR2)-expressing RSC PV interneurons were optically stimulated with 470 nm blue light in PV-Cre or 5XFAD/PV-Cre mice, while whole-cell voltage-clamp recordings were made from RSC excitatory neurons using borosilicate glass electrode (4– 8 MΩ) filled with internal solution containing (in mM) 115 Cesium methanesulfonate (CsMSF), 8 NaCl, 10 HEPES, 0.3 GTP-NaCl, 4 ATP-Mg, 0.3 EGTA, 5 QX-314, and 10 BAPTA (pH[7.3–7.4 and 280–290[mOsm/L). IPSC was recorded at the holding potential of 10 mV. To analyze the stimulus-response (S-R) curve of PV interneuron-evoked IPSC (Figure 3B, C), a single 470 nm blue light pulse (1 ms) with different stimulation intensities (5, 10, 15, 20, and 30 % of maximal light power (15 mW))^1, 2^ was delivered to ChR2-expressing PV interneurons and the corresponding evoked IPSCs were recorded from RSC excitatory neurons. For the subsequent paired-pulse ratio (PPR) analysis (Figure 3D, E), two 470 nm blue light pulses (1 ms) with stimulation intensity that gave the half-maximal IPSC response in the S-R curve^1, 2^ was delivered at 20 Hz. PPR was calculated by normalizing the amplitude of second evoked-IPSC by that of first evoked-IPSC. All signals were amplified (MultiClamp 700B amplifier, Molecular Devices, USA), low-pass filtered at 10 kHz, and acquired at 5 kHz using the ITC-18 data acquisition interface (HEKA Elektronik, Germany). Igor Pro software (WaveMetrics, USA) was used for generating command signals and acquiring data after which acquired data were then analyzed with custom-made Matlab code.

## QUANTIFICATION AND STATISTICAL ANALYSIS

### Statistics

Data are represented as mean ± SEM. Statistical significance was measured using paired or unpaired Student’s t-test, two-way or one-way ANOVA with *post hoc* Tukey’s test, permutation test, and Chi-squared test of proportions.

## Supplementary Information

**Figure S1.**
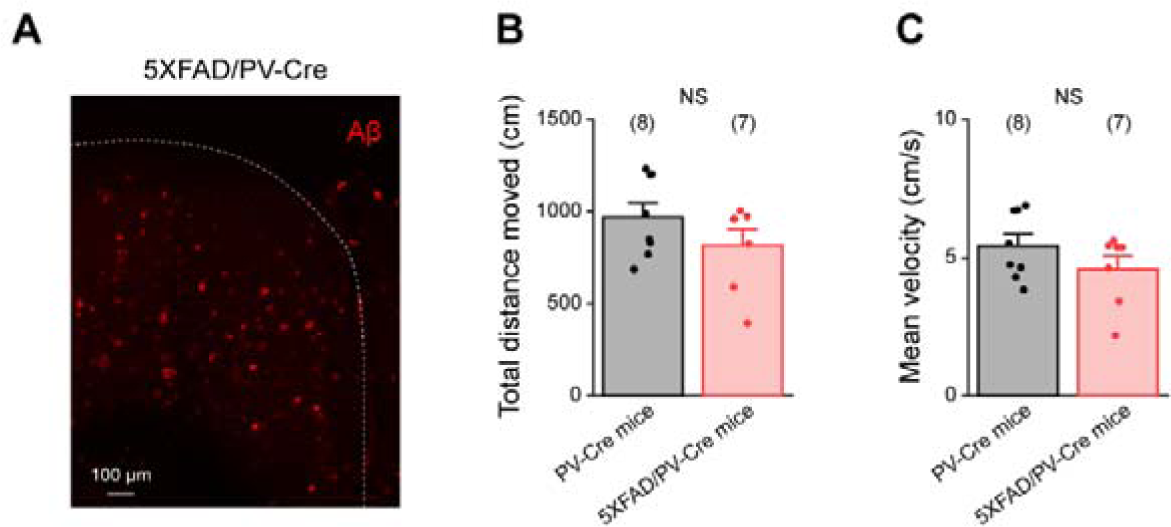
Locomotion analysis of 5XFAD/PV-Cre mice. (**A**) Fluorescent image of beta-amyloid (Aβ) in the granular layer of retrosplenial cortex of 5XFAD/PV-Cre mouse. (**B**, **C**) Averaged total distance travelled (B) and mean velocity (C) of PV-Cre mice and 5XFAD/PV-Cre mice during habituation session. Data are represented as mean ± SEM. n.s.: *p* > 0.05 for unpaired Student’s *t* test (B, C).

**Figure S2.**
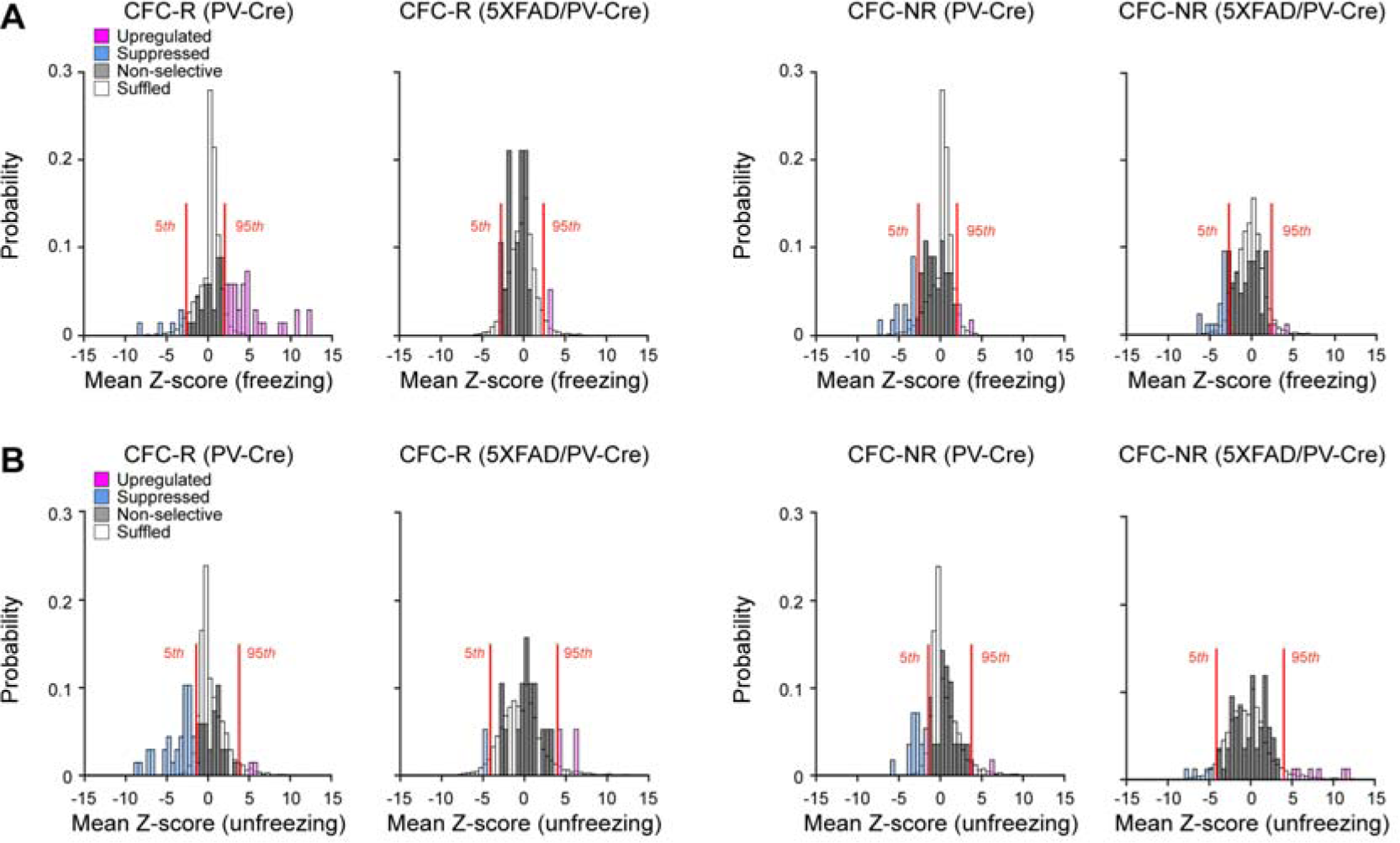
Freezing and unfreezing-selectivity analysis of PV interneuron in the granular retrosplenial cortex. (A, B) Probability distributions of mean Z-score of neuronal activities of CFC-responsive (CFC-R) and CFC non-responsive (CFC-NR) PV interneurons in the granular retrosplenial cortex from PV-Cre mice or 5XFAD/PV-Cre mice during freezing (A) and unfreezing epoch (B) plotted together with the randomly shuffled mean Z-score for comparison (white). Red line: 5th or 95th percentile of randomly shuffled mean Z-score.

**Figure S3.**
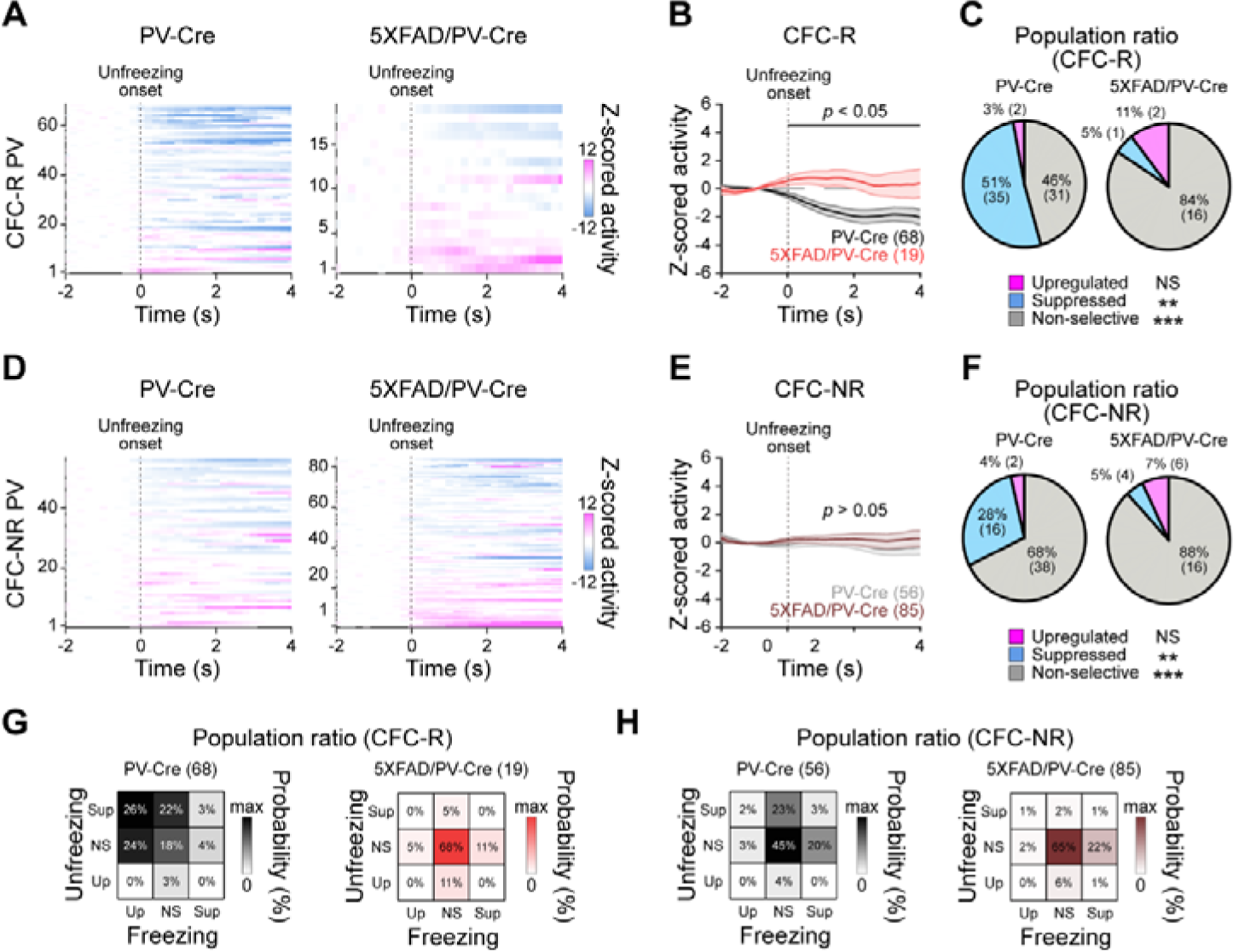
Selective suppression of CFM-encoding PV interneuron ensemble dynamics in the granular retrosplenial cortex during unfreezing epoch of CFM retrieval is disrupted in 5XFAD/PV-Cre mice. (**A**) Z-scored neuronal activity map of CFC-responsive (CFC-R) PV interneurons in the granular retrosplenial cortex at the onset of unfreezing epoch. (**B**) Averaged Z-cored neuronal activities of CFC-R PV interneuron at the onset of unfreezing epoch. Solid line: mean Z-scored activity. Shade: SEM. Black horizontal line: significant differences according to the Permutation test. (**C**) Population ratio of upregulated, suppressed, and non-selective CFC-R PV interneuron according to the mean Z-score at the onset of unfreezing epoch. (**D-F**) Same with (A-C) but for CFC non-responsive (CFC-NR) interneurons from PV-Cre mice or 5XFAD/PV-Cre mice. (**G**, **H**) Population ratio of upregulated (Up), non-selective (NS), and suppressed (Sup) CFC-R (G) and CFC-NR (H) PV interneurons according to the mean Z-score at the onset of freezing and unfreezing epochs. Permutation test (B, E). \*\*\**p* < 0.001, \*\**p* < 0.01, n.s.: *p* > 0.05 for Chi-squared test of proportions (C, F).

**Figure S4.**
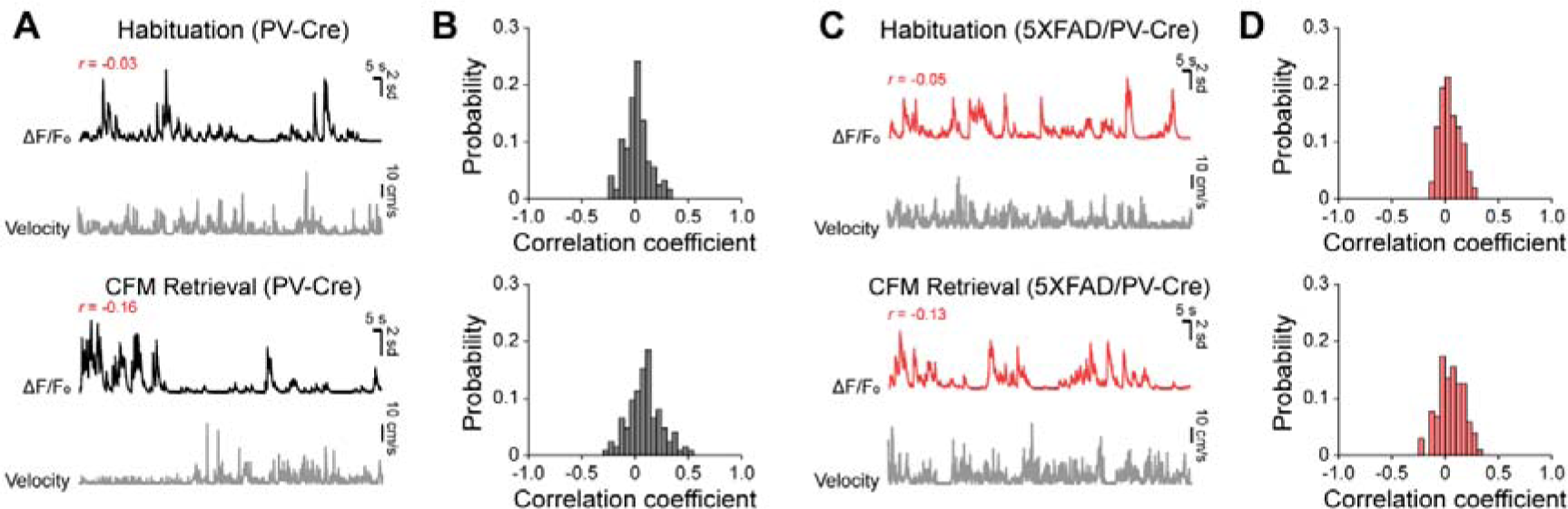
Freezing and unfreezing-selectivity of PV interneurons in the granular retrosplenial cortex is independent from mouse locomotion. (**A**) Representative raw Ca^2+^ signals (△F/F_0_, top) of a single PV interneuron in the granular retrosplenial cortex (RSC) of PC-Cre mouse (black) during habituation and CFM retrieval sessions. Corresponding velocity traces of PV-Cre mice (bottom, gray). Inset: correlation coefficient (*r*) between the representative raw Ca^2+^ signals and the corresponding velocity trace. (**B**) Probability distributions of correlation coefficient between Ca^2+^ signals from all imaged PV interneuron and the velocity traces of corresponding PV-Cre mice during habituation and CFM retrieval sessions. (**C**, **D**) Same with (A, B), but for PV interneuron in RSC of 5XFAD/PV-Cre mice.

